# Intranasal Colonization of Germ-Free Mice with Human Nasal Microbial Communities

**DOI:** 10.64898/2026.04.24.720711

**Authors:** Sudhanshu Shekhar, Anna Michl, Dagmar Šrůtková, Sabina Górska, Fernanda Cristina Petersen, Martin Schwarzer

## Abstract

We present a novel protocol for intranasal colonization of germ-free mice with human nasal microbiota, coupled with an optimized DNA extraction method for murine nasal samples compatible with 16S rRNA sequencing. This protocol may facilitate investigation of the functional roles and causal effects of nasal microbial communities in chronic rhinosinusitis and antimicrobial resistance.

## Introduction

Recent studies have accentuated the therapeutic potential of the nasal microbiota in treating chronic rhinosinusitis and combating antimicrobial resistance [1, 2]. To efficiently reap the benefits of the nasal microbiota, it is critical to gain a better understanding of their functional roles and the causal relationships influencing respiratory host-microbiota interactions. A germ-free (GF) mouse model intranasally colonized with the human nasal microbiota shall serve as a powerful tool for elucidating these functional roles. However, establishing and characterizing such a model poses significant technical challenges, primarily due to difficulties in extracting microbial DNA from murine nasal passages, which contain low microbial biomass and a high proportion of host-derived DNA. Here, we present, for the first time, a protocol for intranasal colonization of GF mice with human nasal microbial communities, along with an optimized method for extracting microbial DNA from murine nasal passages suitable for 16S rRNA next-generation sequencing analysis.

## Methods

### Mice

GF female C57BL/6 mice were maintained under sterile conditions in Trexler-type plastic isolators or an IsoCage Bioexclusion system. Their axenic status was routinely verified by aerobic and anaerobic cultivation of fecal samples and isolator swabs, and Gram staining of fecal smears [3]. Mice were housed under a 12 h light-dark cycle at 22°C and 55% humidity with irradiated bedding (Safe, Rosenberg, Germany) with enrichment nestlets (Ancare, NY, USA), or mouse house (Techniplast, Italy). Mice had *ad libitum* access to autoclaved tap water, and an irradiated (50 kGy) mouse breeding grain-based diet V1124–300 (ssniff Spezialdiäten GmbH, Germany). All procedures were approved by the Committee for the Protection and Use of Experimental Animals of the Institute of Microbiology, Czech Academy of Sciences (approval ID: 153/2024-P).

### Inoculum preparation and transplantation

A nasal swab sample was collected from a healthy adult female donor (approved by the Military Bioethics Committee in Wrocław, Poland; no. 187/21). A sterile swab was inserted into each nostril and transferred into a sterile microcentrifuge tube containing 1 ml of sterile solution (900 µl PBS and 100 µl glycerol). The sample was vortexed for 1 min, aliquoted, and stored at −80°C until use. For colonization of GF mice, a 100 µl aliquot was thawed and diluted 1:3 in sterile PBS. A 10-µl volume of the diluted inoculum was administered intranasally to each mouse using a sterile pipette under a laminar-flow hood at 21 days of age (post-weaning). Following inoculation, mice were housed in sterile IsoCage Bioexclusion cages until the end of the experiment.

### Nasal tissue collection

At days 3 and 7 post-nasal inoculation, mice were anesthetized with isoflurane and euthanized by cervical dislocation. The facial and cranial skin was reflected. A horizontal incision was made below the eyes, followed by a midline vertical incision along the internasal suture, extending from the midpoint between the eyes to the nasal aperture. The nasal bones were elevated with fine forceps to create a bone flap, providing access to the nasal cavity. Intranasal tissues, including the septum, lateral wall, and turbinates, were excised to maximize recovery of colonized bacteria (**Figure 1**). These tissue fragments were transferred to 1.8 ml sterile tubes containing 500 µl PBS and 2.8 mm ceramic beads. Samples were homogenized by vortexing for 1 min and briefly centrifuged at 1,200 rpm. The supernatant was collected and stored at −80°C until further use.

**Figure 1.**
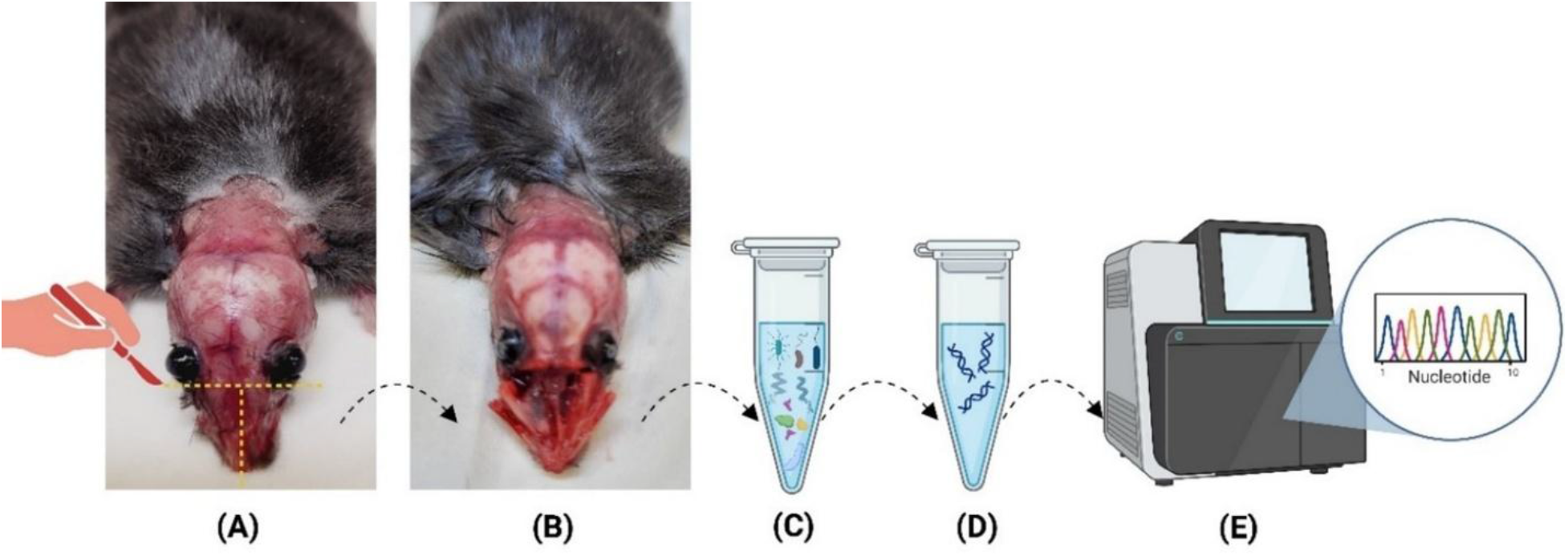
DNA extraction from nasal passages of germ-free mice colonized with the human nasal microbiota for 16S rRNA sequencing. (A) At days 3 and 7 following nasal inoculation, mice were anesthetized and euthanized. The facial and cranial skin was carefully reflected. A horizontal incision was made below the eyes, followed by a midline vertical incision along the internasal suture, extending from the midpoint between the eyes to the nasal aperture. (B) The nasal bones were elevated with fine forceps to create a bone flap, providing access to the nasal cavity. Intranasal tissues, including the septum, lateral wall, and turbinates, were excised. (C) These tissue fragments were transferred to microcentrifuge tubes containing PBS and ceramic beads, followed by thorough vortexing and centrifugation. (D) Microbial DNA was extracted from nasal tissue samples using the Ultra-Deep Microbiome Prep Kit. (E) Extracted DNA was subjected to 16S rRNA gene sequencing.

### Microbial DNA extraction

Microbial DNA was extracted from mouse nasal tissue and human nasal swab samples using the Ultra-Deep Microbiome Prep Kit (Molzym, Germany) following the manufacturer’s instructions. This kit enriches microbial DNA by selectively lysing host DNA while preserving microbial DNA. DNA concentrations were measured using a NanoDrop spectrophotometer (Thermo Fisher Scientific, USA). Extracted DNA was stored at −20°C until shipment to SEQme (Prague, Czech Republic) for 16S rRNA sequencing. Subsequent PCR amplification, sequencing, and data analysis were performed as previously described [4].

## Results

Microbial DNA was extracted from the nasal tissues of GF mice following intranasal inoculation with the human nasal microbiota. On day 3 post-inoculation, DNA yields from the two sampled mice were 7.33 ng/µl and 3.93 ng/µl; on day 7, yields from the remaining two mice were 3.22 ng/µl and 18.03 ng/µl. In contrast, nasal tissues from two uninoculated GF control mice, and an elution buffer negative control, yielded DNA concentrations below the detection limit. 16S rRNA sequencing of DNA recovered from inoculated GF mice demonstrated successful colonization by multiple bacterial taxa. Across both day 3 and day 7 time points, *Lactobacillaceae* were the most abundant family, followed by *Enterococcaceae, Staphylococcaceae*, and *Bacillaceae* (**Figure 2**).

**Figure 2.**
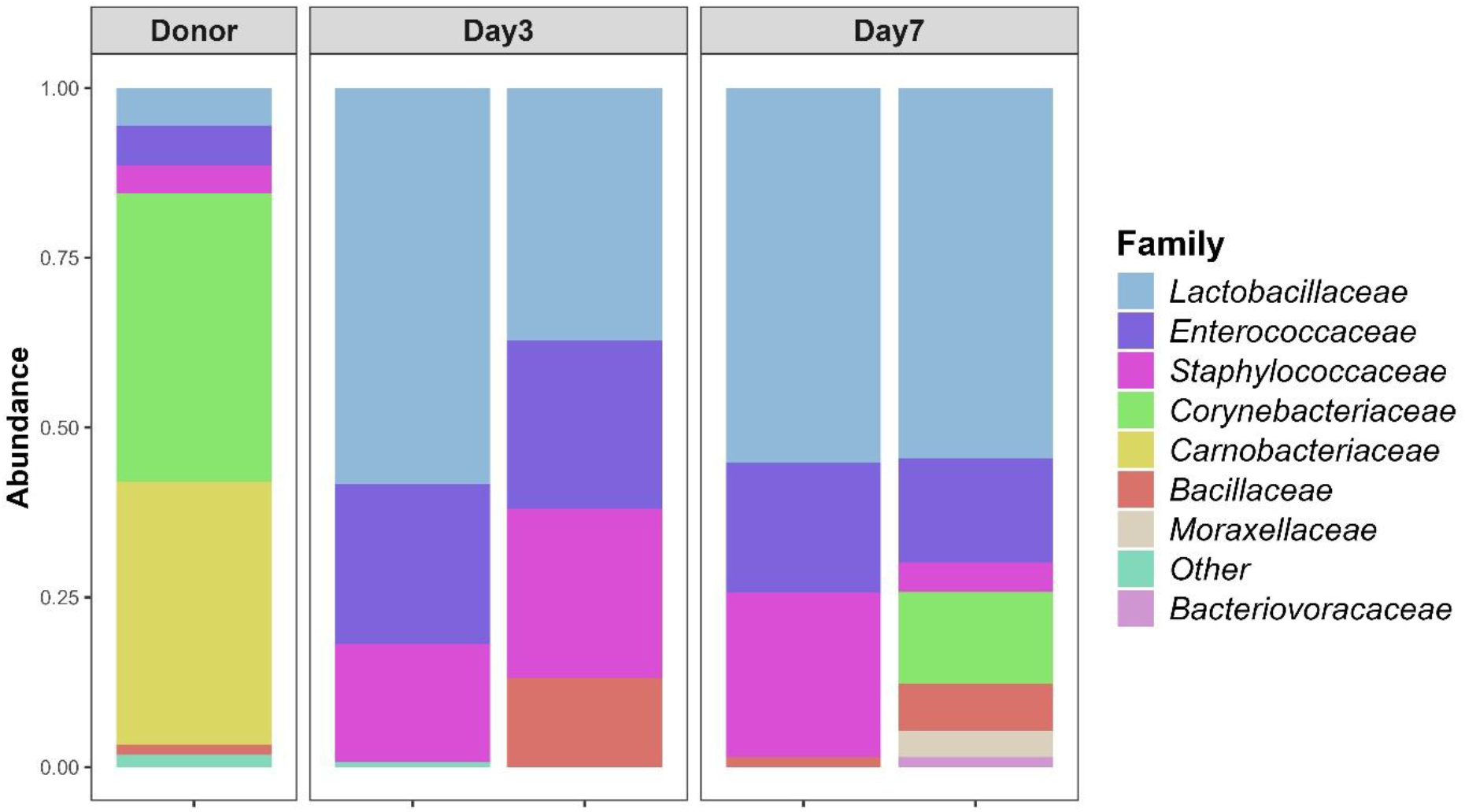
Intranasal colonization of the human nasal microbiota into GF mice. GF C57BL/6 female mice (n = 4; weaning age, postnatal day 21) were intranasally inoculated with the human nasal microbiota. Two mice were euthanized at day 3 and two at day 7 post-inoculation, and nasal tissues were aseptically excised. Microbial DNA isolated from mouse nasal tissues was analyzed by 16S rRNA gene sequencing. The stacked bar plot depicts family-level taxonomic composition of the nasal microbiota, with bars representing the mean relative abundance of taxa within the human donor as well as each mouse sample. Contaminant taxa found in controls were removed during the analysis.

## Discussion

We successfully transferred the human nasal microbiota to GF mice, showing that several microbial taxa can colonize the murine nasal tract. We also established an effective method for isolating microbial DNA from mouse nasal tissues for 16S rRNA sequencing. While the human donor sample contained multiple dominant bacterial families, including *Cornobacteriaceae, Corynebacteriaceae, Staphylococcaceae, Enterococcaceae, Lactobacillaceae*, and *Bacillaceae*, consistent colonization in recipients was observed mainly for *Lactobacillaceae, Enterococcaceae, Staphylococcaceae*, and *Bacillaceae*. These include taxa linked to respiratory homeostasis and antimicrobial resistance, underscoring this method’s utility despite the expected host-specific adaptation.

We isolated microbial DNA directly from nasal tissues rather than from nasal washes, providing a more accurate representation of the resident microbiota, as previously shown [5]. Although yields varied, consistent detection of amplifiable DNA indicates persistence of transplanted communities for at least one week. Overall, this gnotobiotic model offers a novel platform for investigating the human nasal microbiota and host-microbe interactions in the respiratory tract.

## Acknowledgements

We would like to acknowledge support from the Ministry of Education, Youth and Sports of the Czech Republic - Talking Microbes: understanding microbial interactions within the One Health framework (<u>CZ.02.01.01/00/22_008/0004597</u>), the Czech Science Foundation (grant 26-19874L), and the National Science Centre (OPUS 28+LAP/Weave no. UMO-2024/55/I/NZ7/02334). In addition, we thank Šárka Maisnerová for her excellent technical support with the germ-free mouse experiments.

